# Environmental differentiation during virtual navigation in the absence of entorhinal grid-like signals

**DOI:** 10.64898/2026.11.23.768598

**Authors:** Paul F. Hill, Sabina Srokova, Arne D. Ekstrom

**Affiliations:** University of Arizona Psychology Department; McKnight Brain Institute

**Keywords:** Spatial Navigation, Entorhinal Cortex, Parahippocampal Place Area, Grid-Like Signals, Pattern Separation, Scene Perception

## Abstract

The neural mechanisms supporting the encoding and differentiation of distinct spatial environments, particularly the contributions of environmental boundaries and scene-related information, remain unclear. Here, participants encoded two spatial environments during passive virtual navigation: an open outdoor environment without visible boundaries and an indoor environment defined by rectangular boundaries. Based on preregistered hypotheses, we examined whether 6-fold and 4-fold grid-like modulation of the fMRI BOLD signal in entorhinal cortex contributed to the representation of these environments, alongside analyses of environment-specific differentiation in distributed patterns of neural activity. We found no reliable evidence of 6-fold or 4-fold grid-like modulation in either entorhinal cortex or a more anatomically constrained medial entorhinal cortex. Similarly, voxel-wise activity patterns within entorhinal cortex did not reliably differentiate indoor and outdoor environments. In contrast, exploratory analyses revealed significant differences in both univariate BOLD responses and voxel-wise pattern similarity within the scene-selective parahippocampal place area (PPA), with stronger responses and greater differentiation during indoor navigation. We consider these findings in the context of previous studies reporting both positive and null effects of N-fold grid-like modulation in entorhinal cortex, highlighting potential methodological and experimental factors that may influence the detection of these signals. Together, our findings suggest that scene-selective posterior parahippocampal regions may provide a more robust fMRI signature of indoor versus outdoor environmental differentiation under passive navigation conditions, while underscoring the need to better characterize the boundary conditions under which entorhinal grid-like modulation can be reliably detected.

---

Grid cells are neurons in the medial entorhinal cortex of the rodent that fire at multiple locations to form a hexagonal lattice across an environment (Hafting et al., 2005). The metric theory of grid cell coding argues that this periodic spatial organization provides a coordinate system for representing self-location and continuously updating position as an animal moves through space. Grid fields show realignment with movement to different spatial environments such that the orientation, scale, and symmetry of the grid signal are influenced by factors such as environmental boundaries and geometry (Krupic et al., 2015) (Fyhn et al., 2007.

Grid cells within local neuronal ensembles tend to share a common orientation, providing a potential basis for detecting macroscopic grid-like modulation of the fMRI BOLD signal as a function of movement direction. Doeller et al. (2010 demonstrated evidence of 6-fold modulation of the entorhinal fMRI BOLD signal as a function of movement direction and speed during navigation in a virtual environment. Subsequent work with fMRI showed that grid-like modulation of the BOLD signal may be anchored to salient environmental boundaries and barriers (He & Brown, 2019; Julian et al., 2018). In one study, introducing hairpin barriers within a virtual environment resulted in 4-fold modulation of the fMRI BOLD signal, in contrast to the canonical 6-fold modulation observed during open-field navigation (He & Brown, 2019). This finding is broadly consistent with rodent evidence that compartmentalized environments can reorganize grid representations into repeating local submaps (Derdikman et al., 2009).

Together, these findings suggest that entorhinal spatial representations are shaped by environmental context and geometry. This raises a broader question for human fMRI studies: whether rotational symmetry in the fMRI BOLD signal contributes to the encoding of distinct spatial environments, and whether different environments are represented through shifts in grid orientation, changes in rotational symmetry, or other forms of spatial coding, such as spatial pattern separation.

Building on the rodent findings, there are at least two ways in which grid-like modulation of the entorhinal fMRI BOLD signal may differentiate between distinct environments. First, different environments may preserve a common 6-fold rotational symmetry while eliciting different preferred grid orientations, such that the primary axes of the 6-fold modulation reflected in the BOLD signal realign across environments according to salient landmarks, boundaries, or other environmental cues. Alternatively, environmental geometry may alter the rotational symmetry expressed in the entorhinal BOLD signal itself. Rather than simply rotating a common 6-fold representation, different environments may elicit different forms of periodic modulation depending on their geometric structure. In the present study, this would predict classic 6-fold modulation when navigating an open outdoor environment, but greater 4-fold modulation when navigating an indoor environment, where movement is constrained by walls, corners, and dominant north-south/east-west axes (e.g., He & Brown, 2019; Krupic et al., 2015).

Beyond grid-like modulation, distinct environments may also be differentiated through other spatial and scene-based representational mechanisms. Posteromedial regions of the entorhinal cortex have been implicated in representing spatial information and resolving spatial mnemonic interference, potentially providing differentiated spatial inputs to hippocampal circuitry supporting pattern separation (Maass et al., 2015; Navarro Schröder et al., 2015; Reagh & Yassa, 2014). Alternatively, environmental identity may be represented through distributed patterns of voxel-wise activity in posterior scene-selective regions, including posterior parahippocampal cortex. These regions are critical for scene perception and form key components of the broader spatial-navigation network, representing information about scene identity, spatial layout, boundaries, and other structural features of the local environment even in the absence of environment-specific differences in grid-like modulation (Çukur et al., 2016; Ekstrom et al., 2017; R. A. Epstein & Baker, 2019; R. Epstein & Kanwisher, 1998; Kamps et al., 2016; Park & Chun, 2009).

In the present study, participants completed a passive navigation task in which they were guided along fixed trajectories through two distinct environments, encountering trial-unique items along the routes. Following each set of navigation blocks, participants completed a spatial source memory task in which they indicated the environment in which each item had been encountered. We conducted preregistered analyses (Hill & Ekstrom, 2025) to examine whether perceptually distinct indoor and outdoor environments are differentiated through the organization of grid-like modulation or through across-voxel representational differentiation in the entorhinal cortex, and whether the strength of these signals predicts subsequent memory for spatial context. We supplemented these preregistered analyses with a series of exploratory analyses in the entorhinal cortex and scene-selective posterior parahippocampal cortex (PHC) to further characterize environment-related differences in neural activity and representational organization.

## Methods

### Participants

Twenty-eight young adult participants were recruited from the University of Arizona and surrounding communities to participate. Two participants withdrew before completing the study due to claustrophobia and mild motion sickness from the VR task. Data from two additional participants was excluded due to chance level behavioral performance. The final sample comprised 24 young adults (18-40 yrs.; M = 23 yrs.; SD = 4.6 yrs.; 13 females). All participants gave informed consent in accordance with the University of Arizona Institutional Review Board and were compensated at the rate of $20 per hour. All participants were right-handed, had normal or corrected to normal vision, normal or corrected to normal hearing, and reported no history of neurological or psychiatric conditions.

### Virtual Environments

Participants completed the passive navigation task (described below) in each of two virtual environments built in Unity 3D (Unity Technologies, San Francisco, CA). Each environment comprised a navigable space of approximately 60 x 60 virtual meters. Participants were required to freely navigate each environment on a laptop computer prior to the scanning session to familiarize themselves with the layout and visual features of each environment.

The two virtual environments were rendered as distinct indoor and outdoor scenes. The outdoor environment reflected an open desert scene bounded by a circular wall. Distal landmarks located beyond the boundary wall (e.g., palm trees, mountains, cacti, and a windmill) provided directional information that could be used to infer heading. The indoor environment reflected a large rectangular warehouse. The back wall and the large garage-style door indicated the north-south axis of the environment, while the two side walls contained distinct landmarks (Arizona flag and Callahan sign, respectively) that indicated the east-west axis. Both environments included a textured ground surface to convey optic flow during translation and rotation events in the passive navigation task.

**Figure 1.**
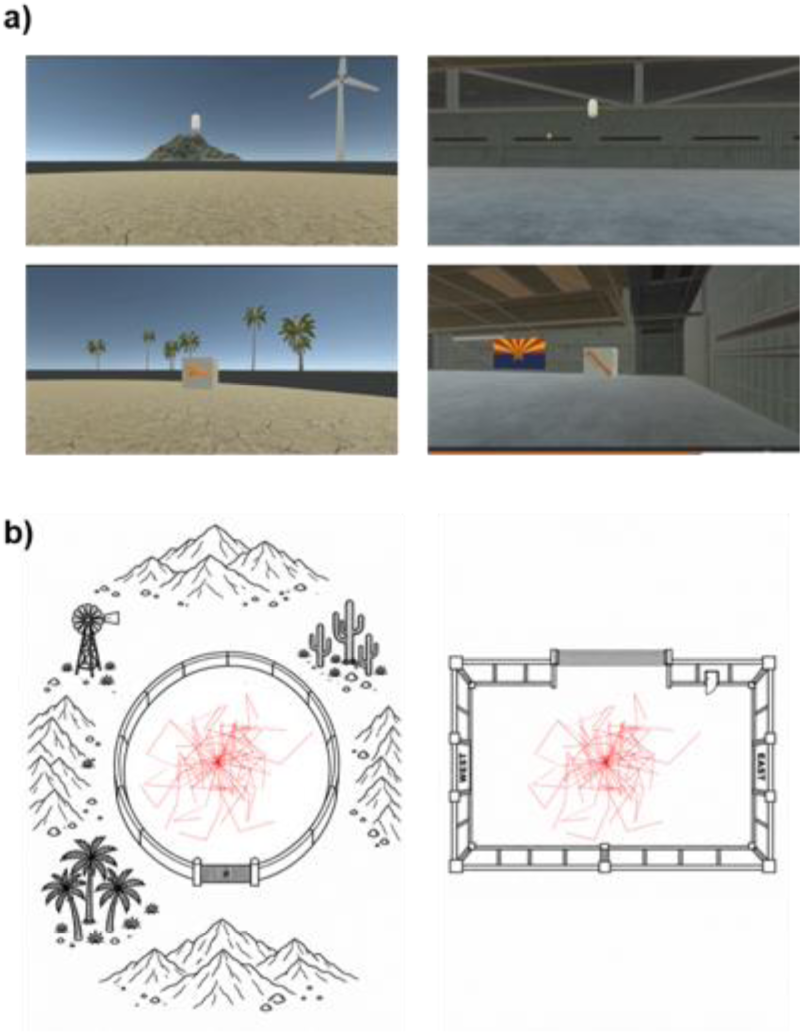
Environmental Task Schematic. (A) First-person views of the individual waypoints (top row) and target item (bottom row) in the outdoor (left column) and indoor (right column) environments. (B) Top-down views illustrating the navigation trajectories, which were matched across the outdoor and indoor environments.

### Passive Navigation Task

Each environment contained a set of 36 trial-unique objects positioned at predetermined locations (72 objects total). On each trial, participants began at a fixed start location in the center of the navigable space and were passively guided along a path consisting of three linear translation segments separated by two rotation events. Participants traveled from the start position to a visually presented waypoint, were automatically rotated to face the direction of the next waypoint and then translated to a second waypoint before a final rotation and translation to the trial-unique object. Upon reaching the final object, which was visually presented on a three-dimensional cube, participants rated how likely the object belonged within the environment (1 = Not Likely, 2 = Somewhat Likely, 3 = Very Likely). These ratings were included to maintain task engagement and were not analyzed further.

The duration of each translation event varied between 2 and 8 seconds. The location, length, and trajectory of each path were matched across environments such that translational and rotational events were evenly distributed across all movement directions and identical between environments. This design ensured uniform sampling of the full 360° directional space, which is essential for computing directional tuning and assessing *N*-fold rotational symmetry.

### Source Memory Task

The 72 objects encountered during the passive navigation task and 24 novel objects not previously encountered were presented individually during a subsequent recognition memory test. Each test item required one or two memory judgments. Participants first indicated whether an item had been encountered during the study phase by making an “Old” or “New” response. Participants were instructed to refrain from guessing and to respond “Old” only when confident that the item had been encountered during the passive navigation task.

Items receiving a “New” response were followed by a white fixation cross for the remainder of the trial. For items receiving an “Old” response, participants additionally indicated whether the object had originally been learned in the indoor or outdoor environment. A “Don’t Know” response option was provided to discourage guessing. All responses were made using a scanner-compatible button box operated with the index, middle, or ring fingers of the right hand.

### Procedures

The experiment consisted of two study-test cycles completed inside the scanner. Each cycle consisted of two consecutive runs of the passive navigation task followed by two consecutive scanning runs of the source memory task. Each run comprised a mixture of indoor and outdoor trials. A brief break separated the study and test phases as participants were reminded of the task instructions. Each passive navigation run included 18 trials split evenly between the indoor and outdoor environments, resulting in 54 total translation events per run (i.e., three translation segments/trial x 18 trials). Each source memory run included 24 trials, including nine outdoor objects, nine indoor objects, and six novel objects.

### MRI Data Acquisition and Preprocessing

Functional and anatomical images were acquired with a 3 T Seimens Skyra MRI scanner equipped with a 32-channel receiver head coil. Functional images were acquired using a T2*-weighted, blood-oxygen-level-dependent echo-planar imaging (EPI) sequence (in-plane parallel imaging acceleration factor = 2, flip angle = 82°, 128 × 128 matrix, FOV = 234 × 234 mm, multiband factor = 3, repetition time [TR] = 2000 ms, echo time [TE] = 30 ms, and right-to-left phase encoding). EPI volumes consisted of 75 slices with 1.8-mm isotropic voxels and no interslice gap. Slices were acquired perpendicular to the long axis of the hippocampus.

Immediately following the second and final study-test cycle, a dual-echo gradient-echo field map was acquired to correct for susceptibility-induced distortions in the functional images (TR = 467 ms, TE1 = 4.92 ms, TE2 = 7.38 ms, flip angle = 60°, 74 × 74 matrix, 42 slices, approximately 3-mm isotropic voxels, no interslice gap). T1-weighted anatomical images were acquired using a three-dimensional magnetization-prepared rapid gradient-echo (MPRAGE) pulse sequence (TR = 2100 ms, TE = 2.33 ms, TI = 1100 ms, flip angle = 12°, 1-mm isotropic voxels, 176 slices sagittal acquisition). High-resolution anatomical images of the hippocampus, entorhinal cortex, and surrounding medial temporal lobe cortex were acquired using a 2D T2-weighted turbo spin-echo sequence (field of view = 215 × 215 mm, matrix = 512 × 512, voxel size = 0.42 × 0.42 × 2.0 mm, repetition time [TR] = 3500 ms, echo time [TE] = 36 ms, flip angle = 120°, 36 slices, echo train length = 24, bandwidth = 222 Hz/pixel).

MRI data were preprocessed using fMRIPrep 25.2.5, based on Nipype 1.10.0. T1-weighted images were corrected for intensity nonuniformity, skull stripped, segmented into gray matter, white matter, and cerebrospinal fluid, and processed with FreeSurfer for cortical surface reconstruction. For each BOLD run, fMRIPrep generated a reference image, estimated head motion using MCFLIRT, applied fieldmap-based susceptibility distortion correction, and coregistered the BOLD reference to the T1-weighted image using boundary-based registration. Confound estimates included framewise displacement, DVARS, global signals, and anatomical and temporal CompCor components. Motion parameters and nuisance regressors were written to the corresponding confounds files, and frames exceeding 0.5 mm framewise displacement or 1.5 standardized DVARS were flagged as motion outliers. Functional data were resampled using a single interpolation step combining the relevant motion, distortion-correction, and spatial transforms. Each of the following analyses was performed on functional data retained in each participant’s native space and spatially smoothed with a 4.5-mm full-width at half-maximum Gaussian kernel.

### ROI Definitions

High-resolution T2 images were automatically segmented in ITK-SNAP using the Princeton Young Adult 3T ASHS Atlas (Aly & Turk-Browne, 2015). All entorhinal masks were resliced to EPI space and manually inspected to ensure accurate anatomical coverage. We further manually divided the entorhinal cortex into medial (MEC) and lateral (LEC) components.

Because rodent grid cells are most commonly recorded in dorsomedial entorhinal cortex, corresponding broadly to the posterior-medial extent of the human entorhinal cortex, we reasoned that grid-like signals might be strongest within the MEC ROI. MEC was defined in reference to the uncal apex, beginning four slices anterior to the uncal apex and including all slices posterior to that point. LEC was defined by identifying the first appearance of the hippocampal head, moving two slices posterior, and including all slices anterior to that point. These boundaries were intended to provide a practical separation between MEC and LEC rather than to imply discrete anatomical borders, given evidence that entorhinal organization varies along overlapping anatomical and functional gradients (Maass et al., 2015; Navarro Schröder et al., 2015; Syversen et al., 2021). The separation was therefore designed to minimize inclusion of the transition zone between putative MEC and LEC territories.

Left and right posterior parahippocampal cortex (PHC) ROIs were identified from the Schaefer 1000-parcel, 17-network atlas in MNI space. Because the atlas does not contain a single parcel corresponding precisely to the canonical parahippocampal place area, adjacent parcels spanning posterior parahippocampal and ventral occipitotemporal cortex were transformed separately into each participant’s native T1-weighted anatomical space and subsequently resliced to the mean functional EPI image.

**Figure 2.**
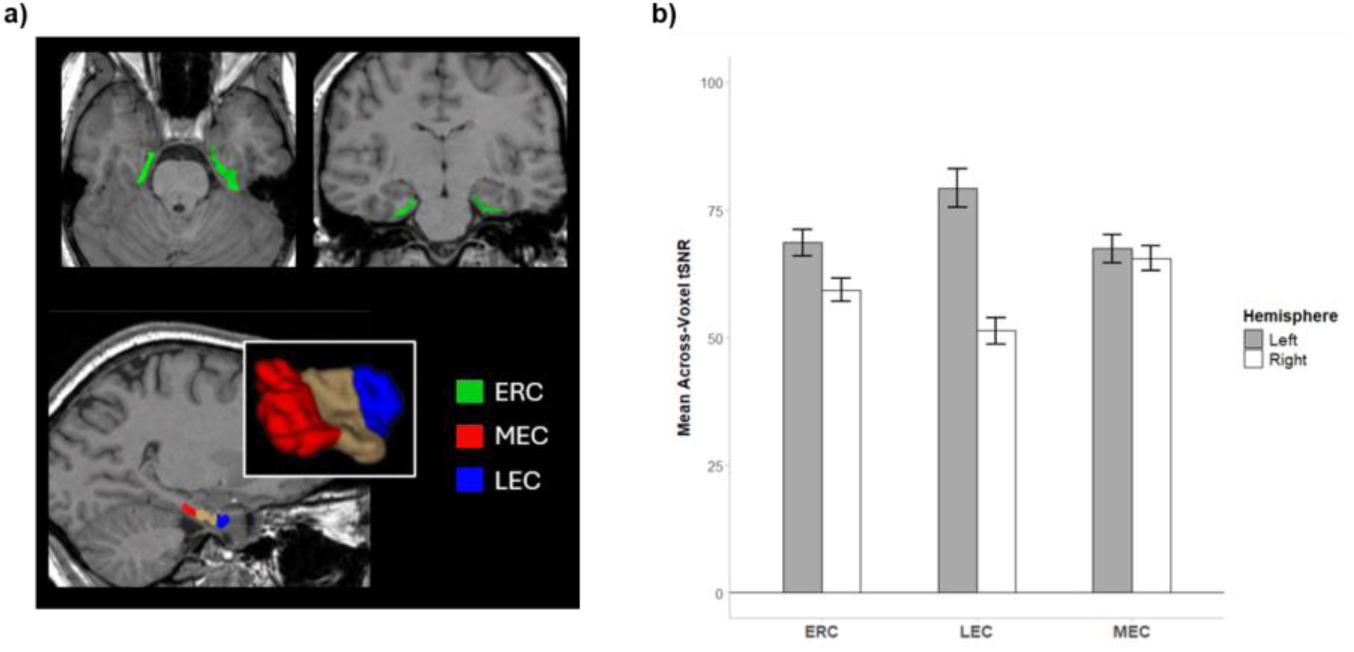
Entorhinal ROI Definitions and Signal Quality. (**A**) Top row: Axial and coronal slices illustrating bilateral entorhinal cortex ROIs (top row). The bottom row illustrates the subdivision of the entorhinal cortex into anterolateral (LEC) and posteromedial (MEC) ROIs, defined using anatomical landmarks corresponding to the hippocampal head and uncal apex, respectively. (B) Temporal signal-to-noise ratio (tSNR) computed from preprocessed functional data within the entorhinal ROIs.

Temporal signal-to-noise ratio (tSNR) was calculated from the preprocessed functional data retained in each participant’s native space and spatially smoothed with a 4.5-mm full-width at half-maximum Gaussian kernel to assess functional signal quality within the entorhinal ROIs. For each functional run, tSNR was calculated separately for each voxel as the temporal mean BOLD signal divided by its temporal standard deviation. Only voxels falling within both the anatomical ROI and the first-level GLM analysis mask were included. Voxelwise tSNR values were then averaged across voxels within each ROI, and the resulting ROI-level estimates were averaged across runs to obtain a participant-level measure of signal quality. Across participants, mean tSNR was generally robust across entorhinal ROIs. Although there was no main effect of ROI, *F*(1.09, 25.18) = 0.80, *p* = .391, tSNR was significantly higher in the left than right hemisphere, *F*(1, 23) = 32.06, *p* < .001, and this effect varied by ROI, as indicated by a significant ROI × hemisphere interaction, *F*(1.08, 24.94) = 45.11, *p* < .001. Follow-up comparisons indicated significantly higher tSNR in left than right ERC (*p* = .003) and LEC (*p* < .001), whereas left and right MEC did not differ (*p* = .966). The lowest tSNR values were observed in right LEC.

### MRI Analyses

#### N-fold BOLD Modulation Analysis

We assessed 4-fold and 6-fold modulation of the fMRI BOLD signal using the Grid Code Analysis Toolbox (GridCAT; Stangl et al., 2017) implemented in MATLAB 2024a. Passive navigation trials were divided into independent estimation and test datasets separately for the indoor and outdoor environments. Each trial contained three translation segments, all of which served as events of interest; intervening rotation events were not included in the model. Across all trials, the first two translation segments were assigned to the estimation dataset, whereas the third and final translation segment was assigned to the independent test dataset. Grid orientation was therefore estimated using the first two translation periods across trials and subsequently tested using the final translation period across trials.

Estimation trials were fit with a first general linear model (GLM1) to estimate preferred grid orientations within each ROI. Under a grid-like coding scheme, BOLD activity is expected to vary systematically with movement direction, following either 6-fold (60°) or 4-fold (90°) rotational symmetry. To identify this directional modulation, GLM1 included two parametric regressors defined as sin(nθ) and cos(nθ), where θ represents the direction of movement within the virtual environment and n corresponds to the rotational symmetry being tested (n = 6 or 4). Separate beta estimates were obtained for each regressor at the voxel level and subsequently averaged across voxels within each ROI. These averaged estimates were then used to calculate the preferred grid orientation for each ROI as φ = atan2(βsin, βcos)/n.

A second GLM (GLM2) was used to model the held-out test trials according to their alignment with the mean grid orientation identified by GLM1. Grid-like modulation was modeled continuously using a parametric modulator ranging from +1 for movement perfectly aligned with the preferred grid orientation to −1 for movement maximally misaligned with that orientation. For each test event, alignment was quantified as cos[*n*(*α* − *θ*)], where *α*was the movement direction, *θ*was the mean grid orientation estimated in GLM1, and *n* corresponded to the symmetry being tested (6 or 4). For 6-fold symmetry, perfectly aligned directions occurred at *θ*, *θ* + 60^∘^, *θ* + 120^∘^, *θ* + 180^∘^, *θ* + 240^∘^, and *θ* + 300^∘^, whereas maximally misaligned directions occurred 30° from these axes, at *θ* + 30^∘^, *θ* + 90^∘^, *θ* + 150^∘^, *θ* + 210^∘^, *θ* + 270^∘^, and *θ* + 330^∘^. For 4-fold symmetry, perfectly aligned directions occurred at *θ*, *θ* + 90^∘^, *θ* + 180^∘^, and *θ* + 270^∘^, whereas maximally misaligned directions occurred at *θ* + 45^∘^, *θ* + 135^∘^, *θ* + 225^∘^, and *θ* + 315^∘^.

#### Grid Stability Analyses

The magnitude of N-fold modulation is influenced by the stability of estimated grid orientations across voxels within an ROI (i.e., spatial stability) and within voxels across scanner runs (i.e., temporal stability). A Rayleigh’s test for non-uniformity of circular data was used to assess whether estimated grid orientations are uniformly distributed across voxels within an ROI, or whether grid orientations across voxels are clustered around a common orientation. Higher Rayleigh Z scores indicate greater spatial stability. Temporal stability was computed by comparing each voxel’s grid orientation across each of the respective scanner runs. A voxel was classified as temporally stable when its estimated orientation differed across runs by no more than half the aligned-misaligned angular separation, corresponding to ± 15° for 6-fold symmetry and ± 22.5° for 4-fold symmetry. Temporal stability was quantified as the average percentage of stable voxels across all pairwise run comparisons. Because 50% stability is expected by chance, temporal stability scores significantly greater than 50% indicate reliable stability across runs.

#### Permutation-based test of N-fold modulation amplitude

Although cross-validation provides a sensitive test of whether an ROI’s preferred orientation generalizes to held-out data, it does not directly assess whether BOLD activity exhibits N-fold rotational symmetry in the first place. This is especially relevant as the preferred orientation can be estimated even when the underlying parameter estimates from the sine and cosine regressors are near zero. For that reason, we also performed a phase-independent permutation analysis that quantified the magnitude of N-fold modulation across the full dataset, irrespective of the ROI’s preferred orientation. Because this approach estimates N-fold symmetry using the full dataset, it places less stringent demands on orientation stability than cross-validation.

For each subject and ROI, directional modulation was modeled using sine and cosine parametric regressors at 6-fold or 4-fold symmetry, analogous to the GLM1 analysis described above. Because the sine and cosine parameter estimates from this GLM jointly encode the periodic response, modulation amplitude for the observed data was computed as *A=√(β^2^_cos_ + β^2^_sin_).* We next generated a subject-level null distribution by repeatedly shuffling movement direction within each run and environment. Instead of re-estimating the full GLM for each permutation, which can introduce nuisance estimation variability in the null distribution, we extracted the filtering (*K*) and whitening (*W*) operators from the observed GLM and held them constant across all permutations. The BOLD time series were first pre-multiplied by the observed *K* and *W*. For each permutation, the shuffled directional regressors were used to specify a new design matrix, which was subjected to the same *K* and *W* transformation. Voxel wise parameter estimates and the resultant modulation amplitude were obtained by multiplying the pseudoinverse of the transformed design matrix with the filtered and whitened BOLD data.

At the group level, permutation-specific modulation amplitudes were averages across subjects to obtain a group level null distribution. The group mean for the observed amplitude was then compared with this distribution using a one-sided permutation test, with significance defined as the proportion of null observations greater than or equal to the observed group mean.

#### Multivoxel Pattern Similarity Analyses

As a complementary analysis, we examined multivoxel patterns of activity within the same entorhinal cortex ROIs using pattern similarity analyses that did not depend on periodic modulation of mean fMRI BOLD amplitude. Analyses were performed separately for indoor and outdoor trials. For each translation event, we extracted both the mean BOLD response within each ROI and the corresponding multivoxel activity pattern. Thus, the pattern similarity analyses could be interpreted alongside conventional differences in response magnitude, allowing us to determine whether effects reflected changes in overall BOLD amplitude, voxel-wise representational structure, or both. We calculated pairwise Pearson correlations between trials within the same environment, and correlation coefficients were Fisher *z*-transformed prior to analysis. To minimize similarity attributable to temporal autocorrelation and other run-specific effects, correlations were calculated only between trials occurring in different scanning runs.

Our first analysis tested whether multivoxel activity patterns exhibited representational similarity consistent with 6-fold or 4-fold rotational symmetry. For the 6-fold analysis, trial pairs were classified as within-symmetry when their movement headings differed by no more than ±15° after accounting for 60° periodicity. Thus, headings separated by approximately 0°, 60°, 120°, 180°, 240°, or 300° were treated as occupying equivalent positions within the hypothesized 6-fold representation. Trial pairs falling outside this range were classified as between-symmetry. For the 4-fold analysis, the same procedure was applied using 90° periodicity, with trial pairs classified as within-symmetry when their headings differed by no more than ±22.5° after accounting for 90° periodicity. Mean Fisher *z*-transformed similarity was then calculated separately for within- and between-symmetry trial pairs within each environment, with greater within-than between-symmetry similarity taken as evidence that multivoxel activity patterns reflected the corresponding rotational similarity.

Next, we performed a heading-direction analysis designed to capture similarity associated with absolute movement direction rather than rotational symmetry. Analyses were again conducted separately for indoor and outdoor trials, with pairwise correlations restricted to trials from different scanning runs. Trial pairs were classified as having similar headings when their movement directions differed by no more than ±15° or ±22.5°, corresponding to the angular tolerances used in the 6-fold and 4-fold analyses, respectively. Unlike the symmetry analyses, no modulo transformation was applied, such that headings were considered similar only when they were close in absolute directional space. Mean Fisher *z*-transformed correlations were then calculated for similar- and dissimilar-heading trial pairs. Greater similarity for trials with similar headings was interpreted as evidence for a general heading-direction signal, potentially reflecting shared visual or directional information associated with movement in the same direction.

Finally, we performed an indoor vs. outdoor similarity analysis to determine whether multivoxel activity patterns within entorhinal cortex differentiate between the two environments. For each indoor trial, within-environment similarity was defined as its mean Fisher *z*-transformed correlation with other indoor trials, whereas between-environment similarity was defined as its mean correlation with outdoor trials. The same procedure was then repeated for outdoor trials, such that within-environment similarity reflected correlations with other outdoor trials and between-environment similarity reflected correlations with indoor trials. As in the preceding analyses, correlations were restricted to trials occurring in different scanning runs.

Greater within-than between-environment similarity was interpreted as evidence that voxel-wise patterns of entorhinal activity differentiated the indoor and outdoor environments.

**Figure 3.**
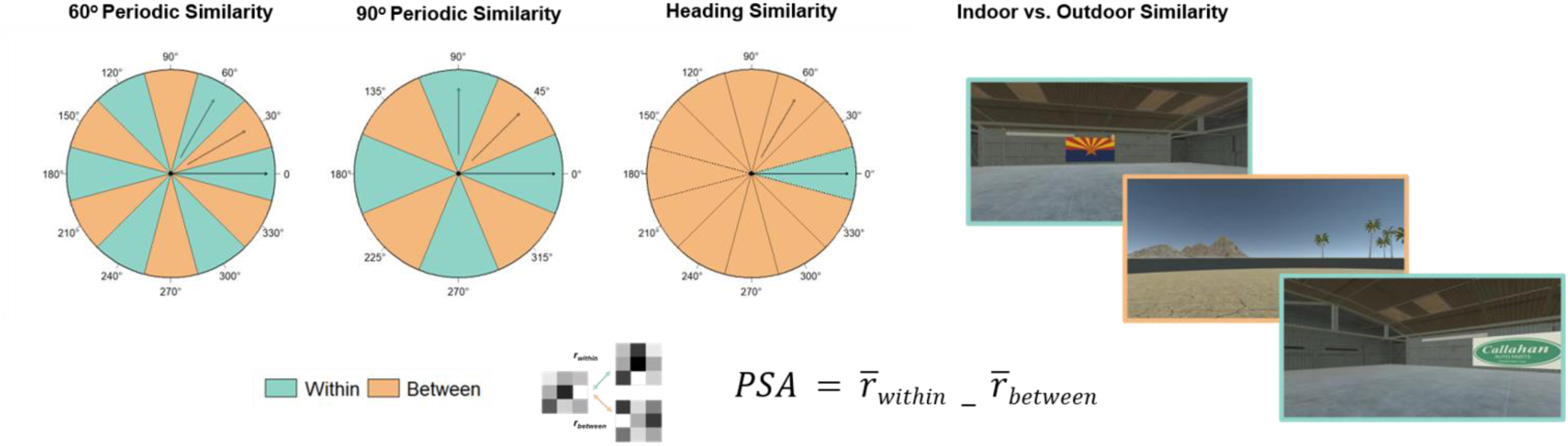
Schematic of voxel-wise pattern similarity analyses. Top Left: Polar plots illustrating the classification of event pairs as within (green) or between (orange) conditions for computing directionally modulated pattern similarity. Top Right: First-person views illustrating the indoor and outdoor environments used to assess environment-specific differentiation. In both analyses, the pattern similarity metric was quantified as the difference between within- and between-condition voxel-wise pattern similarity.

## Results

### Source Memory Performance

Source memory performance was calculated separately for indoor and outdoor items as the proportion of correctly identified source judgments among successfully recognized items (i.e., item hits). Overall, source memory performance was high for both indoor (M = .78, SD = 12) and outdoor (M = .76, SD = .14) items. Performance on the source memory task did not significantly differ between the indoor and outdoor environments (*t*(44) = 0.60, *p* = .146).

### Weak to no evidence for entorhinal grid-like modulation in the fMRI BOLD signal

To test our preregistered hypotheses regarding environment-dependent differences in grid-like modulation of the entorhinal fMRI BOLD signal, we performed separate 2 × 2 ANOVAs with factors of hemisphere (left, right) and environment (indoor, outdoor) for 4-fold and 6-fold rotational symmetry. Within the entire ERC ROI, the analysis of 6-fold symmetry revealed no significant main effects of environment (*F*(1, 23) = 3.26, *p* = .084, η²p = .124, BF10 = 1.77) or hemisphere (*F*(1, 23) = 1.04, *p* = .319, η²p = .043, BF10 = 0.36), and no significant environment × hemisphere interaction (*F*(1, 23) = 0.07, *p* = .791, η²p = .003, BF10 = 0.32). We note that Bayes Factor scores provide anecdotal evidence in favor of an effect of environment on 6-fold symmetry in the entorhinal cortex. To further characterize this effect and determine whether 6-fold modulation differed reliably from zero, we conducted follow-up one-sample *t*-tests (two-tailed) against a null value of zero. These analyses were collapsed across hemisphere given the absence of significant hemisphere effects. Six-fold modulation did not differ significantly from zero in either the indoor environment, *t*(23) = −1.06, *p* = .301, or the outdoor environment, *t*(23) = 1.83, *p* = .081. For consistency with prior studies of grid-like modulation, which have commonly reported one-tailed tests, we also note that a one-tailed test indicated significant 6-fold modulation in the outdoor environment, *t*(23) = 1.83, *p* = .040, whereas modulation in the indoor environment did not differ from zero, *t*(23) = −1.06, *p* = .850. We interpret this result cautiously and report it primarily for comparability with the existing literature.

An identical 2 × 2 ANOVA of 4-fold symmetry revealed no significant main effects of environment (*F*(1, 23) = 1.34, *p* = .260, η²p = .055, BF10 = 0.39) or hemisphere (*F*(1, 23) = 3.25, *p* = .085, η²p = .124, BF10 = 1.36), and no significant environment × hemisphere interaction (*F*(1, 23) = 1.87, *p* = .185, η²p = .075, BF10 = 0.58). Follow-up one-sample *t*-tests (two-tailed) against a null value of zero confirmed that 4-fold modulation did not significantly differ from zero in either the indoor environment, *t*(23) = −1.32, *p* = .200, or the outdoor environment, *t*(23) = 0.23, *p* = .818. These conclusions were unchanged when one-tailed one-sample *t*-tests were performed for the indoor, *t*(23) = −1.32, *p* = .900, and outdoor environments, *t*(23) = 0.23, *p* = .409.

Within the MEC ROI, the analysis of 6-fold symmetry revealed a significant main effect of environment (*F*(1, 23) = 5.16, *p* = .033, η²p = .183, BF10 = 3.25), driven by greater 6-fold modulation in the outdoor compared with the indoor environment. Neither the main effect of hemisphere (*F*(1, 23) = 0.12, *p* = .728, η²p = .005, BF10 = 0.23) nor the hemisphere × environment interaction (*F*(1, 23) = 0.69, *p* = .414, η²p = .029, FB10 = 0.38) was significant. Follow-up one-sample *t*-tests (two-tailed), collapsed across hemisphere, indicated that 6-fold MEC modulation did not reliably differ from zero in either the indoor environment, *t*(23) = −1.77, *p* = .091, or the outdoor environment, *t*(23) = 1.69, *p* = .104. For consistency with prior studies of grid-like modulation, we also conducted exploratory one-tailed tests. Six-fold modulation in the outdoor environment approached, but did not reach, the conventional threshold for statistical significance, *t*(23) = 1.69, *p* = .052, whereas modulation in the indoor environment did not differ from zero, *t*(23) = −1.77, *p* = .955.

**Figure 4.**
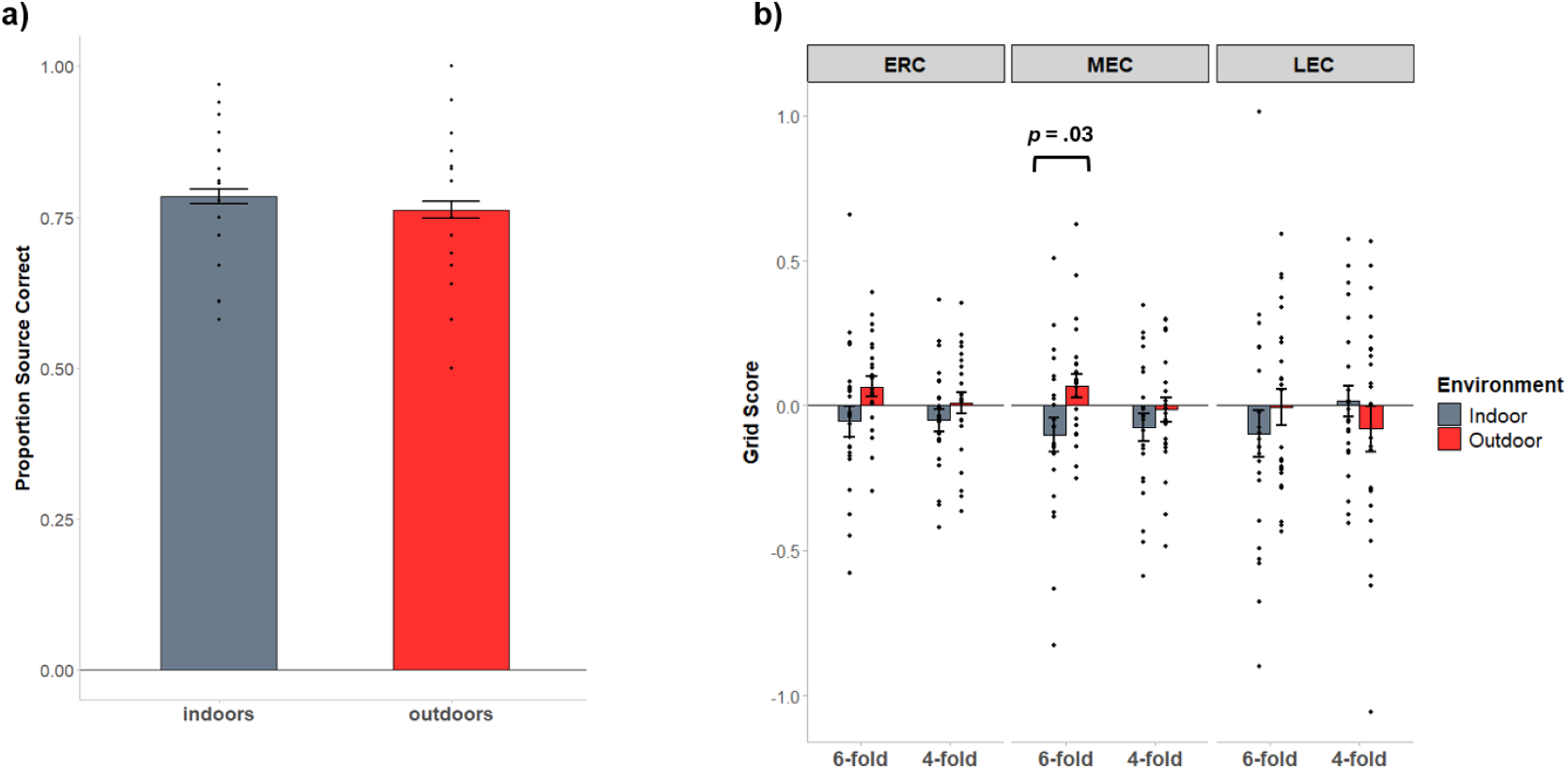
Behavioral Performance and N-Fold BOLD Modulation. (A) Source memory performance was significantly above chance and did not differ between indoor and outdoor environments. (B) Neither 6-fold nor 4-fold BOLD modulation differed significantly from zero in any entorhinal ROI or environment. However, a significant main effect of environment was observed for 6-fold modulation in the MEC.

An identical 2 × 2 ANOVA of 4-fold symmetry within the MEC revealed no significant main effects of environment (*F*(1, 23) = 0.99, *p* = .329, η²p = .041,BF10 = 0.318) or hemisphere, (*F*(1, 23) = 0.16, *p* = .693, η²p = .007, BF10 = 0.24) and no significant environment × hemisphere interaction (*F*(1, 23) = 0.006, *p* = .940, η²p < .001, BF10 = 0.26). Follow-up one-sample *t*-tests (two-tailed), collapsed across hemisphere, confirmed that 4-fold modulation did not significantly differ from zero in either the indoor environment, *t*(23) = −1.56, *p* = .133, or the outdoor environment, *t*(23) = −0.34, *p* = .739. These conclusions were unchanged in exploratory one-tailed tests for the indoor, *t*(23) = −1.56, *p* = .934, and outdoor environments, *t*(23) = −0.34, *p* = .630.

Within the LEC ROI, 6-fold symmetry showed no significant effects of environment, hemisphere, or their interaction (all *ps* > .35, all BF10 < 0.37), and modulation did not differ from zero in either environment. Four-fold symmetry likewise showed no environment effect or environment × hemisphere interaction, although there was a weak main effect of hemisphere (*F*(1, 23) = 4.33, *p* = .049, η²p = .159, BF10 = 2.01).

### Spatial and Temporal Stability

We computed estimates of spatial stability separately for the indoor and outdoor environments and for 4-fold and 6-fold symmetry. As shown in Figure 5, Rayleigh *Z* scores were generally high across conditions, with a majority of participants exhibiting spatially coherent, non-uniform grid orientations. For 6-fold symmetry, Rayleigh *Z* scores did not differ significantly by environment, hemisphere, or their interaction within ERC (all *ps* > .11) or MEC (all *ps* > .18). Within LEC, there was a significant main effect of hemisphere, *F*(1, 23) = 20.30, *p* < .001, η²G = .205, reflecting greater spatial coherence in right than left LEC, but no significant effect of environment or environment × hemisphere interaction (both *ps* > .19).

**Figure 5.**
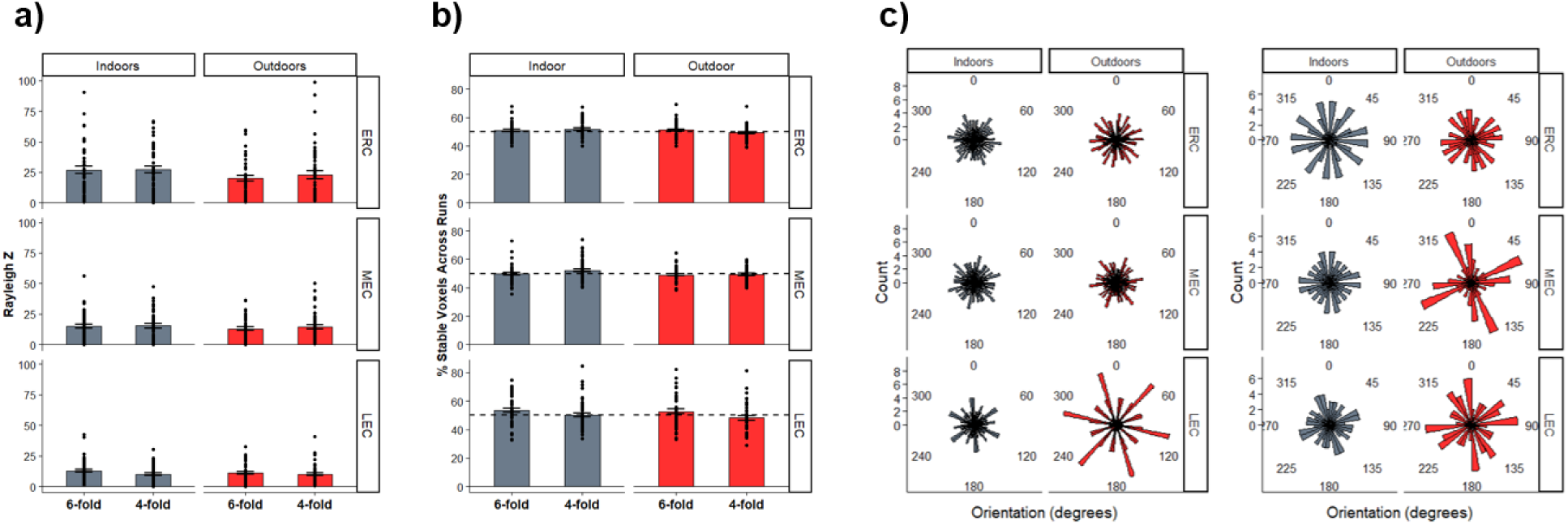
Spatial and Temporal Stability of N-Fold Modulation. (A) Rayleigh Z-scores reflecting the spatial clustering of preferred grid orientations across voxels within each entorhinal ROI. Preferred orientations were generally nonuniformly distributed, suggesting some degree of spatial clustering. (B) Percentage of voxels exhibiting stable preferred grid orientations across scanner runs, which did not significantly exceed chance levels. (C) Polar histograms illustrating the distribution of preferred grid orientations across participants. Rayleigh tests indicated no significant clustering, providing no evidence that preferred grid orientations were systematically aligned with common visual landmarks or environmental geometry across participants.

A similar pattern was observed for 4-fold symmetry. Rayleigh *Z* scores did not differ significantly by environment, hemisphere, or their interaction within ERC (all *ps* > .17) or MEC (all *ps* > .24). Within LEC, there was again a significant main effect of hemisphere, *F*(1, 23) = 9.77, *p* = .005, η²G = .099, reflecting greater spatial coherence in right than left LEC, with no significant effect of environment or environment × hemisphere interaction (both *ps* > .29).

To assess whether preferred orientations were also aligned across participants, we submitted each participant’s mean voxel-wise, across-run grid orientation to a group-level Rayleigh test of circular uniformity separately for each ROI, environment, and symmetry model. Preferred orientations were not significantly aligned across participants in ERC, MEC, or LEC for either 4-fold or 6-fold symmetry (all *ps* > .05). Thus, although individual participants generally exhibited coherent across-voxel orientations within ROIs, the orientation of this coherence was not reliably shared across participants.

We also estimated temporal stability separately for the indoor and outdoor environments and for the 4-fold and 6-fold symmetry conditions. For 6-fold symmetry, the percentage of stable voxels did not differ significantly by hemisphere, environment, or their interaction within ERC, MEC, or LEC (all *ps* > .15). Temporal stability also did not significantly exceed the 50% chance level in any 6-fold condition. For 4-fold symmetry, the percentage of stable voxels likewise did not differ significantly by hemisphere, environment, or their interaction within ERC, MEC, or LEC (all *ps* > .09). Relative to chance, stability was generally not significantly greater than 50%; the sole exception was left ERC in the indoor environment, where stability was modestly above chance, *M* = 52.6%, *t*(23) = 2.27, *p* = .033. Left MEC indoors showed a similar trend, *M* = 53.0%, *t*(23) = 2.02, *p* = .056.

### Permutation-based test of N-fold modulation amplitude

Permutation tests did not provide evidence for 4-fold or 6-fold modulation amplitudes relative to the shuffled null. Indeed, as illustrated in Fig 6, 6-fold modulation in ERC, MEC, and LEC resulted in p_perm_ > 0.274, while 4-fold modulation was p_perm_ > 0.633. The results were unchanged when evaluating the lower tail of the permutation distribution; no observed modulation amplitudes were significantly lower than expected under the shuffled-direction null. Therefore, although the cross-validation analysis yielded weak evidence for positive 6-fold modulation in the ERC under a one-tailed test, this effect was not accompanied by an above-null chance in overall 6-fold modulation amplitude. This discrepancy raises the possibility that the weak evidence for 6-fold modulation observed in the cross-validation analysis may reflect a false positive, particularly given that the effect was not supported by permutation-based analyses using an empirical null distribution.

**Figure 6.**
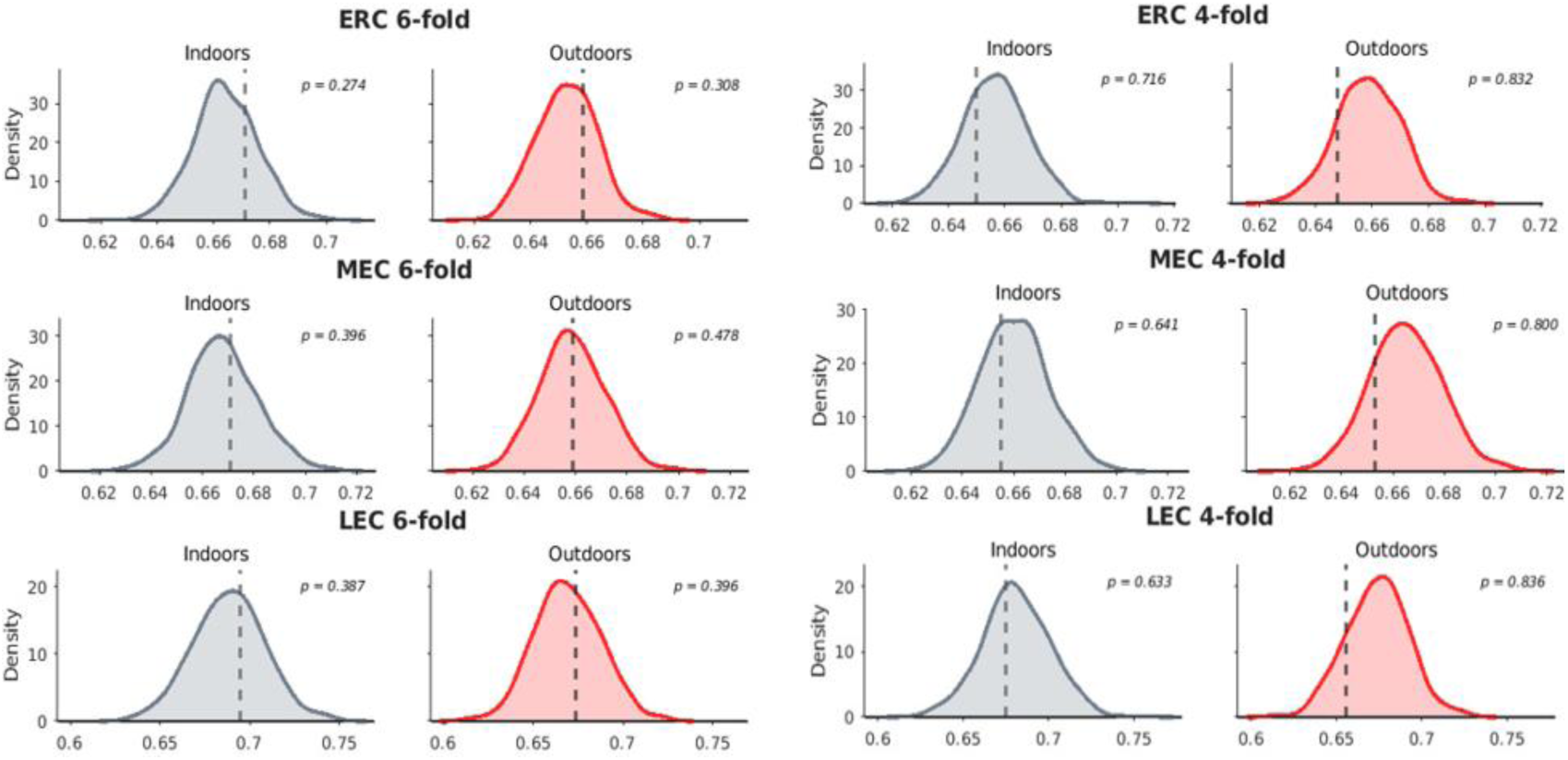
Permutation-based tests of modulation amplitude. Kernel density estimates reflect group level permutation null distributions, separately for indoor and outdoor environments. Dashed vertical line indicates observed group mean.

### Environmental and Directional Pattern Similarity in the Entorhinal ROIs

We next used multivoxel pattern similarity analyses to test our preregistered hypothesis that voxel-wise activity patterns within the entorhinal ROIs (ERC, MEC, and LEC) reliably differentiate between indoor and outdoor environments. Pattern similarity analyses in ERC revealed no significant main effects of hemisphere (F(1, 23) = 0.02, p = .877, η²p = .001, BF10 = 0.22) or environment (F(1, 23) = 0.02, p = .876, η²p = .001, BF10 = 0.21), and no hemisphere × environment interaction (F(1, 23) = 0.52, p = .480, η²p = .022, BF10 = 0.37). Similarly, no significant effects of hemisphere, environment, or their interaction were observed in MEC (all ps ≥ .072, all BF10s < 0.67) or LEC (all ps ≥ .131, all BF10s < 0.87). Follow-up two-tailed one-sample t-tests indicated that pattern similarity did not differ significantly from zero for indoor or outdoor events in any entorhinal ROI (all ps ≥ .136).

Univariate mean BOLD responses within the entorhinal ROIs similarly showed little evidence of systematic environmental modulation. In ERC, there were no significant main effects of hemisphere (*F*(1, 23) = 3.13, *p* = .090, η²p = .120, BF10 = 0.49), or environment (*F*(1, 23) = 1.22, *p* = .281, η²p = .050,BF10 = 0.53), and no hemisphere × environment interaction (*F*(1, 23) = 0.63, *p* = .436, η²p = .027, BF10 = 0.35). Likewise, no significant effects were observed in MEC (all *ps* ≥ .224, all BF10s < 0.94) or LEC (all *ps* ≥ .184, all BF10s < 0.56).

Follow-up one-sample *t*-tests indicated that mean BOLD responses did not differ significantly from zero for indoor or outdoor events in either hemisphere for ERC or MEC (all *ps* ≥ .099). In LEC, mean BOLD response was significantly below zero for outdoor events in the left hemisphere, *t*(23) = −2.88, *p* = .009, whereas the remaining LEC conditions did not differ significantly from zero (all *ps* ≥ .084).

We next supplemented the preregistered indoor versus outdoor differentiation analyses with a series of exploratory directional similarity analyses to examine whether voxel-wise activity patterns exhibited rotational symmetry as a function of movement direction. We first examined whether voxel-wise pattern similarity exhibited 6-fold (60°) rotational symmetry by grouping pairwise differences in movement direction into 30° bins (±15°) centered at 0°, 60°, 120°, 180°, 240°, and 300°. Pattern similarity was then compared between trials with angular differences falling within these bins and trials with angular differences falling outside these bins. The main effects of environment, hemisphere, and the environment x hemisphere interactions were not significant in any of the entorhinal ROIs. Likewise, none of the condition-specific pattern similarity estimated significantly differed from zero (all ps > .184).

We next examined whether voxel-wise pattern similarity exhibited 4-fold (90°) rotational symmetry. Within the ERC, the main effect of environment was not statistically significant (*F*(1, 23) = 3.26, p = .084, η²p = .124). The Bayesian analysis, however, did provide moderate evidence favoring a main effect of environment (BF10 = 4.93). This pattern was characterized by greater 4-fold rotational similarity for indoor trials compared to outdoor trials. Neither the main effect of hemisphere (F(1, 23) = .15, p = .702, η²p = .006, BF10 = 0.21) nor the environment x hemisphere interaction (F(1, 23) = 0.00, p = .983, η²p < .001, BF10 = 0.29) was significant. Pattern similarity during outdoor trials was significantly less than zero, t(23) = −2.27, p = .033.

A similar pattern of 4-fold symmetry was also evident in the MEC, for which the main effect of environment was not statistically significant (*F*(1, 23) = 3.14, *p* = .090, η²p = .210), although the Bayesian analysis provided weak evidence favoring a main effect of environment (BF₁₀ = 2.36). Neither the main effect of hemisphere (*F*(1, 23) = 1.14, *p* = .296, η²p = .047, BF₁₀ = 0.34) nor the environment × hemisphere interaction (*F*(1, 23) = 0.34, *p* = .565, η²p = .015, BF₁₀ = 0.29) was statistically significant. Within the LEC, there was a significant main effect of hemisphere (F(1, 23) = 4.56, p = .044, η²p = .166, BF₁₀ = 1.77). The main effect of environment (*F*(1, 23) = 0.05, *p* = .296, η²p = .002, BF₁₀ = 0.22) and the environment x hemisphere interaction were not significant (*F*(1, 23) = 2.06, *p* = .164, η²p = .082, BF₁₀ = 0.53).

In additional exploratory analyses, we examined whether voxel-wise pattern similarity exhibited a direction-specific heading effect by comparing similarity between trials with movement directions differing by no more than 30° (±30°) to trials with angular differences exceeding 30°, consistent with a preference for specific heading directions rather than rotational symmetry. The main effects of environment and the environment x hemisphere interaction were not significant in any of the entorhinal ROIs (all ps > .140, all BF10s > 0.66). A significant main effect of hemisphere was observed in ERC (F(1, 23) = 4.49, p = .045, η²p = .163, BF10 = 1.17) and was slightly beyond the a priori significance threshold in the MEC (F(1, 23) = 4.25, p = .051, η²p = .156, BF10 = 1.11). In both cases, this effect was driven by greater heading similarity in the left hemisphere relative to the right hemisphere. The main effect of hemisphere in the LEC was not significant (F(1, 23) = 1.61, p = .217, η²p = .065, BF10 = 0.57). None of the condition-specific heading-direction pattern similarity estimates differed significantly from zero (all ps > .056).

### Environmental and Directional Pattern Similarity in Posterior Parahippocampal Cortex

Given the absence of indoor-outdoor differentiation in the entorhinal ROIs, we next conducted exploratory analyses to determine whether environmental differentiation was evident in scene-selective regions of the PHC. In contrast to the entorhinal ROIs, pattern similarity analyses in left and right PPC revealed significant main effects of hemisphere, *F*(1, 23) = 5.06, *p* = .034, η²p = .180, BF10 = 0.37), and environment (*F*(1, 23) = 5.88, *p* = .024, η²p = .204, BF10 = 129.55) with no significant hemisphere × environment interaction (*F*(1, 23) = 3.07, *p* = .093, η²p = .118, BF10 = 0.78). Follow-up two-tailed one-sample *t*-tests confirmed significant differentiation for indoor events in both left PHC, *t*(23) = 4.27, *p* = .0003, and right PHC, *t*(23) = 5.05, *p* < .0001, whereas pattern similarity did not significantly differ from zero for outdoor events in either hemisphere (left: *t*(23) = 1.62, *p* = .119; right: *t*(23) = 1.53, *p* = .140).

Mean BOLD responses showed a similar environmental effect, with a significant main effect of environment (*F*(1, 23) = 10.63, *p* = .003, η²p = .316, BF10 = 69.85), but no significant main effect of hemisphere (*F*(1, 23) = 3.34, *p* = .081, η²p = .127, BF10 = 2.50) and no hemisphere × environment interaction (*F*(1, 23) = 0.02, *p* = .899, η²p < .001, BF10 = 0.26). Mean BOLD responses were significantly greater than zero for indoor events in both left PHC, *t*(23) = 2.18, *p* = .040, and right PHC, *t*(23) = 3.14, *p* = .005, but not for outdoor events in either hemisphere (both *ps* ≥ .479).

**Figure 7.**
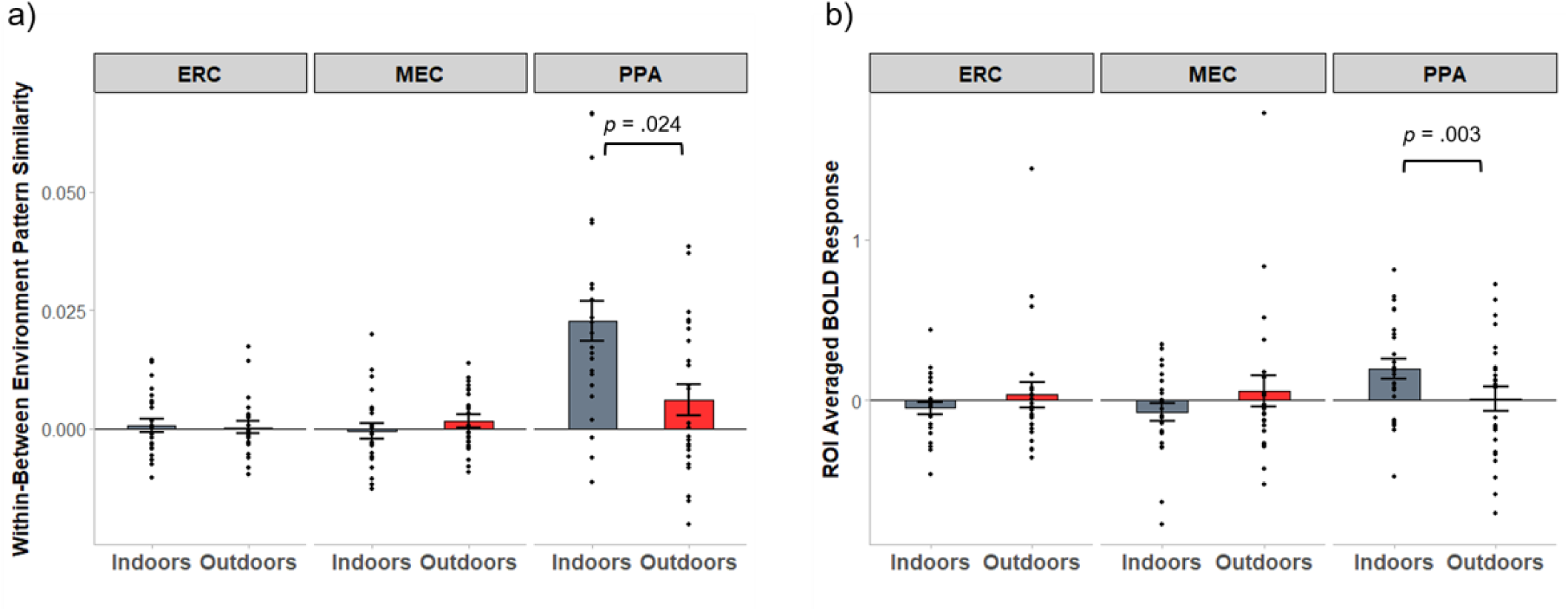
Indoor vs. Outdoor Scene Differentiation. (A) Voxel-wise pattern similarity and (B) ROI-averaged BOLD responses in bilateral ERC, MEC, and PPC. Both analyses revealed significant scene differentiation in the PPC, with greater pattern differentiation and BOLD responses during indoor compared to outdoor navigation.

### Associations of N-Fold Modulation and Pattern Similarity with Spatial Source Memory Performance

Finally, we tested our preregistered hypotheses that individual differences in the magnitude of grid-like modulation and voxel-wise pattern similarity would be associated with subsequent memory for item-context associations. We performed multiple regression analyses with proportion source correct as the dependent variable and the respective fMRI measure, hemisphere, and environment as predictors. These analyses were conducted in the ERC, with corresponding exploratory analyses performed in the scene-selective PHC. We also conducted simple correlations to determine whether the magnitude of each fMRI measure was significantly associated with source memory performance. None of these analyses revealed significant associations between the fMRI measures and subsequent source memory performance (all *ps* > .150).

## Discussion

Across complementary univariate, grid-like modulation, and multivoxel pattern similarity analyses guided by preregistered hypotheses and analytical approaches (Hill & Ekstrom, 2025), we found little evidence that entorhinal cortex reliably differentiated between spatial environments during a passive navigation task. In contrast, scene-selective PHC showed clear environment-specific differentiation effects. Despite the absence of reliable grid-like signals, source memory performance was well above chance in both environments, indicating that participants successfully encoded and subsequently retrieved associations between encountered items and their environmental contexts. Thus, the absence of robust grid-like signals cannot readily be attributed to a general failure to encode information about the respective environments.

Although we observed some evidence of environment-dependent differences in rotational N-fold modulation of the BOLD signal and voxel-wise pattern similarity, these effects were inconsistent across symmetry models and analytical approaches. Within MEC, 6-fold BOLD modulation was significantly greater during outdoor than indoor navigation, but modulation amplitudes did not reliably differ from zero in either environment. Similarly, exploratory pattern similarity analyses revealed greater 4-fold rotational similarity during indoor relative to outdoor navigation within ERC and MEC, although neither environment exhibited significant 4-fold similarity relative to zero. Moreover, the environmental differences observed in 6-fold BOLD modulation were not reproduced in the corresponding pattern similarity analyses, and permutation analyses did not establish modulation amplitudes exceeding chance. Collectively, these findings suggest possible environment-dependent differences in the rotational organization of entorhinal BOLD responses, but provide insufficient evidence for reliable grid-like modulation or consistent effects of environmental context on rotational coding.

Several methodological factors may have limited our ability to detect reliable grid-like modulation in the present study. Participants completed a passive navigation task involving movement along predetermined routes rather than actively controlling navigation. Although grid-like modulation has previously been observed during passive navigation (Kim & Maguire, 2019; Wagner et al., 2023), two recent studies employing similar paradigms also failed to detect reliable 6-fold modulation in the entorhinal cortex (Kransberg et al., 2026; Newton et al., 2023). Notably, both passive and active navigation involve optic flow and a sense of movement, and thus the lack of active movement should not have prevented the emergence of N-aligned grid codes. In addition, findings across studies employing active and passive navigation have been mixed (see Table 1), suggesting that active engagement alone may not reliably determine whether grid-like modulation is observed. Nevertheless, the absence of active navigational decisions, motor planning, and internally generated movement predictions during passive navigation may influence the engagement of grid-related spatial computations and contribute to variability across studies.

**Table 1.** Summary of Prior N-Fold Navigation Task and Imaging Parameters.

| Citation | N-Fold Symmetry | Task | fMRI Slice Orientation |
|---|---|---|---|
| Doeller et al., 2010 | 6-fold | Active Navigation | Not Reported* |
| Horner et al., 2016 | 6-fold | Active Navigation | Oblique, 45º anterior-upward tilt |
| Horner et al., 2016 | 6-fold | Imagined Navigation | Oblique, 45º anterior-upward tilt |
| Stangl et al., 2018 | 6-fold | Active Navigation | Parallel to hippocampal long-axis |
| Kim & Maguire, 2019 | 6-fold | Passive Navigation | Oblique transverse, ~30º tilt |
| Moon et al., 2022 | 6-fold | Active Navigation | Not Reported |
| Wagner et al., 2023 | 6-fold | Observed Navigation | Parallel to hippocampal long-axis |
| Sigismondi et al., 2024 | 6-fold | Imagined Navigation-Sighted Participants | Oblique, 15º relative to AC-PC |
| Sigismondi et al., 2024 | 4-fold | Imagined Navigation-Blind Participants | Oblique, 15º relative to AC-PC |
| Kunz et al., 2015 | 6-fold | Active Navigation | Oblique transverse, ~30º tilt |
| Julian & Doeller, 2021 | 6-fold | Active Navigation | Parallel to hippocampal long-axis |
| He & Brown, 2019 | 6-fold | Active Navigation – Open Field | Parallel to hippocampal long-axis |
| He & Brown, 2019 | 4-fold | Active Navigation – Hairpin Barriers | Parallel to hippocampal long-axis |
| Wagner et al., 2023 | Null | Active Navigation | Parallel to hippocampal long-axis |
| Kransberg et al., 2026 | Null | Passive Navigatin | Not Reported |
| Newton et al., 2023 | Null | Passive Navigation | Parallel to hippocampal long-axis |
| Doeller et al. (2010) cite Weiskopf et al. (2006) for EPI optimization but do not explicitly state EPI slice tilt |  |  |  |

A second consideration could relate to the number of navigation trials available for estimating grid orientation and modulation strength. Dividing trials between indoor and outdoor environments, as well as navigation and retrieval conditions, reduced the data available for estimating environment-specific grid-like signals, potentially increasing variability in orientation estimates and reducing sensitivity to rotationally symmetric modulation. Additionally, indoor and outdoor navigation trials were interspersed within blocks, requiring repeated transitions between distinct spatial environments. Given evidence that grid representations can realign in response to changes in environmental context and geometry (Fyhn et al., 2007), these transitions may have reduced the stability of environment-specific grid orientations across trials.

Finally, differences in fMRI acquisition parameters may have influenced sensitivity to grid-like modulation within the entorhinal cortex, which is particularly susceptible to signal loss and geometric distortion. However, previous studies reporting grid-like modulation have employed varying acquisition orientations, including those parallel and perpendicular to the hippocampal long axis (Table 1), suggesting that acquisition orientation alone is unlikely to account for our null findings. Collectively, these methodological factors may have reduced sensitivity to grid-like modulation, and our null findings should not necessarily be interpreted as evidence for the absence of underlying grid-related spatial representations. Instead, the present results should be seen as adding to the growing number of examples (Table 1) in which 6-fold, and now 4-fold symmetry, are not observed.

Beyond the absence of reliable grid-like modulation, we also found no evidence that distributed patterns of entorhinal activity differentiated between indoor and outdoor environments, despite successful subsequent memory for item-environment associations. This finding is notable given theoretical accounts proposing that posteromedial entorhinal cortex contributes to spatial representations and the resolution of spatial mnemonic interference (Reagh & Yassa, 2014), potentially supporting differentiated representations of distinct environments before information reaches the hippocampus. Our findings, however, do not rule out more fine-grained spatial representations or interference-resolution mechanisms that may not be detectable through voxel-wise pattern similarity analyses.

In contrast to the entorhinal cortex, scene-selective posterior parahippocampal cortex (PHC) showed robust differentiation between indoor and outdoor environments, with significantly greater BOLD responses and more differentiated voxel-wise activity patterns during indoor navigation. These findings are consistent with previous work demonstrating greater PHC activation in response to indoor compared with outdoor scenes (Henderson et al., 2007), as well as evidence that scene-selective regions are particularly sensitive to the geometric properties of visual environments (Epstein & Kanwisher, 1998; Ferrara & Park, 2016; Park et al., 2011).

Specifically, PHC has been shown to respond preferentially to concave compared with convex objects and scene boundaries (Cheng et al., 2021), and to represent geometric properties of the surfaces that define local space (Dillon et al., 2018). Such properties are particularly prominent in indoor environments, where walls, corners, and other structural boundaries constrain the geometry of the surrounding space. Our findings extend this work by demonstrating that environment-specific differences in PHC activity are also evident during dynamic visual navigation through perceptually distinct indoor and outdoor environments, rather than solely during the presentation of static scenes.

Despite the absence of reliable grid-like BOLD modulation in the entorhinal cortex, participants generally demonstrated high levels of subsequent source memory performance, indicating that they successfully encoded and retrieved the environmental contexts in which items were encountered. Nevertheless, none of the examined fMRI measures obtained during the navigation phase were significantly associated with subsequent source memory performance.

Neither the magnitude of grid-like modulation nor the degree of environment-specific pattern differentiation in the entorhinal cortex or PHC reliably predicted individual differences in source memory accuracy. One possible explanation is that relatively high source memory accuracy limited behavioral variability, reducing sensitivity to individual differences in neural activity.

In summary, the present findings add to a growing body of evidence suggesting that grid-like BOLD modulation in the human entorhinal cortex may be less consistently observed across experimental paradigms and contexts than is often assumed. Despite successful source memory for environmental context, we found no reliable evidence of 6-fold modulation or environment-specific differentiation of entorhinal activity during passive navigation, while scene-selective PHC exhibited robust differences in both mean BOLD responses and distributed activity patterns between indoor and outdoor environments. These findings underscore the need to better characterize the experimental and methodological conditions under which grid-like modulation can be reliably detected, including the contributions of active versus passive navigation, environmental geometry, task demands, scanning parameters, and analytical approaches. Given the potential for null findings to be underreported (Ioannidis et al., 2014; Rosenthal, 1979), the existing literature may overestimate the prevalence and reliability of grid-like BOLD modulation across experimental paradigms. Together, systematic comparisons across experimental conditions, analytical methods, and consideration of null results will be important for establishing the boundary conditions of this widely used measure and clarifying its relationship to the underlying neural computations that support human spatial navigation.

## Acknowledgements

Research reported in this publication was supported by the Alzheimer’s Association under award number AARF-22-926755 and the National Institute of Neurological Disease and Stroke under award number R01NS076856. Approximately 10% of the total project costs were financed with federal funds (NINDS) in the amount of $16,666. The remaining 90% of project costs were financed by non-governmental sources (Alzheimer’s Association) in the amount of $150,000. The content is soley the responsibility of the authors and does not necessarily represent the official views of the Alzheimer’s Association or National Institutes of Health.

